# Light intensity affects the coloration and structure of chimeric leaves of *Ananas comosus var. bracteatus*

**DOI:** 10.1101/2020.10.27.356857

**Authors:** Wei Yang, Yuke Lin, Yanbin Xue, Meiqin Mao, Xuzixing Zhou, Hao Hu, Jiawen Liu, Lijun Feng, Huiling Zhang, Jiaheng Luo, Jun Ma

**Affiliations:** College of Landscape Architecture, Sichuan Agricultural University, Chengdu, Sichuan, China

## Abstract

*Ananas comosus var. bracteatus* is an important tropical ornamental plant because of its green/white chimeric leaves. The accumulation of anthocyanin makes the leaf red especially the white margin. However, the leaves lost red color in summer and winter. Light intensity is one of the most important factors affecting leaf color along the season. In order to understand the effects of light intensity on the growth and coloration of the chimeric leaves, *Ananas comosus var. bracteatus* was grown under full sunlight, 50% shade and 75% shade for 75 days to evaluate the content of pigments, the color parameters (value L*, a*, b*) and structural histocytology characteristics of chimeric leaves. The results showed that high irradiance was beneficial to keep the chimeric leaves red. However, prolonged exposure to high irradiance caused light damage, some of the leaves wrinkled and even burned. Shading decreased the content of anthocyanin and increased the content of chlorophyll especially in the white margin of the leaves. Numerous chloroplasts were found in the mesophyll cells of the white margin part of chimeric leaves under shading for 75 days. The increase of chlorophyll content resulted in better growth of plants. In order to balance the growth and ornamental value of the leaves, approximately 50% shade is suggested to be the optimum light irradiance condition for *Ananas comosus var. bracteatus* in summer.

## Introduction

Variegated leaf chimeric plants are important ornamental plants because of their unique colorful characteristics. The high diversity of color in chimeric plants was due to the changes of spatial distribution of pigment types, contents and proportions in the leaves, and this change is the result of the interaction between genetic and environmental factors [1–3]. Climate, soil nutrients, and water has long been understood to be primary factors influencing plant growth, and the light intensity is an important environmental factor [4,5]. In green leaves, chlorophyll dominates in pigments causing a green appearance. Chlorophyll deficiency can cause leaves to turn white or yellow. The light intensity can affect the content of chlorophyll, carotenoid, and anthocyanin in the leaves, leading to variation in leaf color [6]. It has been shown that different plants have different adaptability to light intensity. *Photinia* × *fiaseri* grown under full light was the most densely foliated compared shaded plants [7]. Approximately 67% of natural light (the irradiance intensity of sunlight was about 2500 *μmol* m^2^ s^-1^ at noon during September in Jinhua) was concluded to be the optimum light irradiance condition for *Tetrastigma hemsleyanum* Diels et Gilg [4]. The research conducted by Kong et al [8] showed that 30% and 50% of sunlight (the irradiance intensity of sunlight was about 2000 ± 20 *μmol* photons m^-2^ s^-1^ at noon) were the most suitable growth environment for *Mahonia bodinieri* (Gagnep.) Laferr obtaining higher biomass and could improve the economic value. In addition, Light intensity can tightly regulate color change. *Galax urceolata* leaves had a transition from red to green along with the decrease of light intensity [9]; *Kalamegh* leaf transit its color from green to red from time to time under varying intensities of light and shade [10]. Similarly, Shading was reported to improve foliar color of *Kalmia latifolia* L from yellow-green to dark green in the first year of production [11]. The leaves of both *‘Royal Glissade’* and *‘UF06-1-6’* gradually changed from green to red under different light environment, the anthocyanin content increased while chlorophyll content decrease [12]. And a study in *Arabidopsis thaliana* has shown that light intensity can affect the type of anthocyanin [13]. More importantly, the structural and physiological adaptions of plants enable their survival under different light environmental conditions [14]. Indeed, the adaptions are mainly related to the difference in the quantity and distribution of pigments, external morphology and internal anatomy structure of leaves [15,16]. These changes in anatomical structure can maximize photosynthetic efficiency and keep internal temperatures at optimal levels [17,18].

*Ananas comosus var. bracteatus,* belonging to the family Bromeliaceae, is a herbaceous perennial monocot. It has a high ornamental value all year round due to its green/white chimeric leaves. During spring and autumn, the leaves will appear bright red, which improved a lot in ornamental value of *Ananas comosus var. bracteatus* [19]. However, in summer and winter, the redness of the leaves gradually fades, returning to green/white. How to maintain the red color of the leaves and extend the red color viewing period is an important way to improve the ornamental and economic value of *Ananas comosus var. bracteatus*. We have carried out a lot of studies on the molecular mechanism of chimera traits formation of *Ananas comosus var. bracteatus* [20–24]. The genetic characteristics and physiological properties of other variegated leaf chimeric plants have been systematically studied [25–28]. However, few studies have addressed how chimeric plants response to varying light intensity, and the mechanism under it is still unknown.

The objective of our present study was to determine the optimum light intensity for the growth and high ornamental value of *Ananas comosus var. bracteatus* by investigating the effects of different light intensity on the coloration change, pigments content and proportion and internal anatomy structure of chimeric leaves. Our findings may promote the outdoor application of *Ananas comosus var. bracteatus*, which may prove beneficial to the nursery and landscape industries.

## Materials and methods

### Plant materials and shading treatment

Two year old *Ananas comosus var. bracteatus* plants with red margin chimeric leaves were used in this study. The plants were planted in pots containing a mixture of coconut bran and river sand (1:2). The plants were divided into three groups for shading treatments: natural sun-light condition (CK), 50% shading condition (T1), and 75% shading condition (T2). Shading was accomplished by using one or two layers of commercial black cloth shade, and the treatment lasted for 75 days. The fourth and fifth leaves of the plants were sampled from three plants of each treatment every 15 days for further detection.

### Leaf color parameters

The leaf color parameters were measured by using an automatic color meter. The L*, a*, b* values of the marginal and central parts of the chimeric leaf were measured separately with three replicates.

### Pigment content detection

The marginal and central parts of the fresh leaves were used for pigments detection. The chlorophyll and carotenoid levels were detected using the Holm equation and a method previously described [29]. The anthocyanin content detection is according to the methodology described by Ren et al [25]. Three replications were used for each sample.

### Microscope observation

Leaf samples of each treatment were harvested and sliced by double-sided blades, mounted in water, and viewed immediately. The method of paraffin sections is described in more detail by Liu et al [30]. The observation and photographical documentations of the marginal and central parts of the chimeric leaves were carried out using a photomicroscope (Leica DM1000, Leica Microsystems, Ltd, Germany).

## Results and discussion

### Seasonal cycle of leaf color of *Ananas comosus var. bracteatus*

There are significant differences in the chimeric leaf color of *Ananas comosus var. bracteatus* between four seasons. In spring and autumn, the white edge of the chimeric leaf turns to be red. At this time, the leaves are green/white/red mosaicked which increased a lot the ornamental value of *Ananas comosus var. bracteatus.* (Fig 1A,C). However, in summer and winter, the redness of the leaves gradually fades, only the base part of the leaf edge remains a little red (Fig 1B,D).

**Fig 1.**
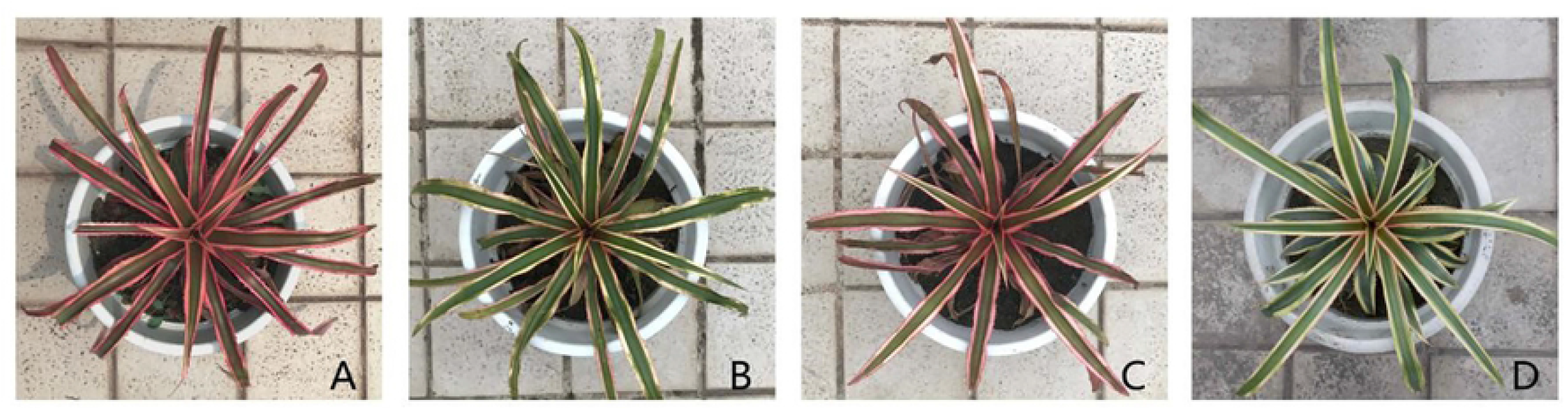
The color change of *Ananas comosus var. bracteatus* leaves along four seasons. spring (A), summer (B), autumn (C), and winter (D).

### Morphologic changes of the leaves under different light conditions

Shading treatment had the significantly effects on *Ananas comosus var. bracteatus* leaf morphology. The morphologic changes of the leaves under different light conditions were shown in figure 2. Plants under 75% and 50% shade grown well throughout the experimental period while sunburn were observed in the leaves of plants under full light. The degree of leaf sunburn increased with the time of exposure. The margin of the chimeric leaves under 75% shade firstly lost the red color to be white (about 45 days) and then turned green along the treatment procedure. The leaf margin under both full light and 50% shade maintained the red color. The sunburn spots existed on the leaves cultured under full light for about 60 days. According to previous studies, light damage was caused by the interaction of excess light and high leaf temperatures and as a consequence affected plant growth [31,32]. Indeed, shading usually reduces plant growth, but some plants, especially many evergreens, benefit from shading in the warm area. For *Euonymus japonica* Hand. -Mazz. ‘Aureo-marginata’, plant growth was optimized with 50% shading [33]. Shading increased the growth of *Rhododendron ×* ‘Pink Ruffles’ [34].

**Fig 2.**
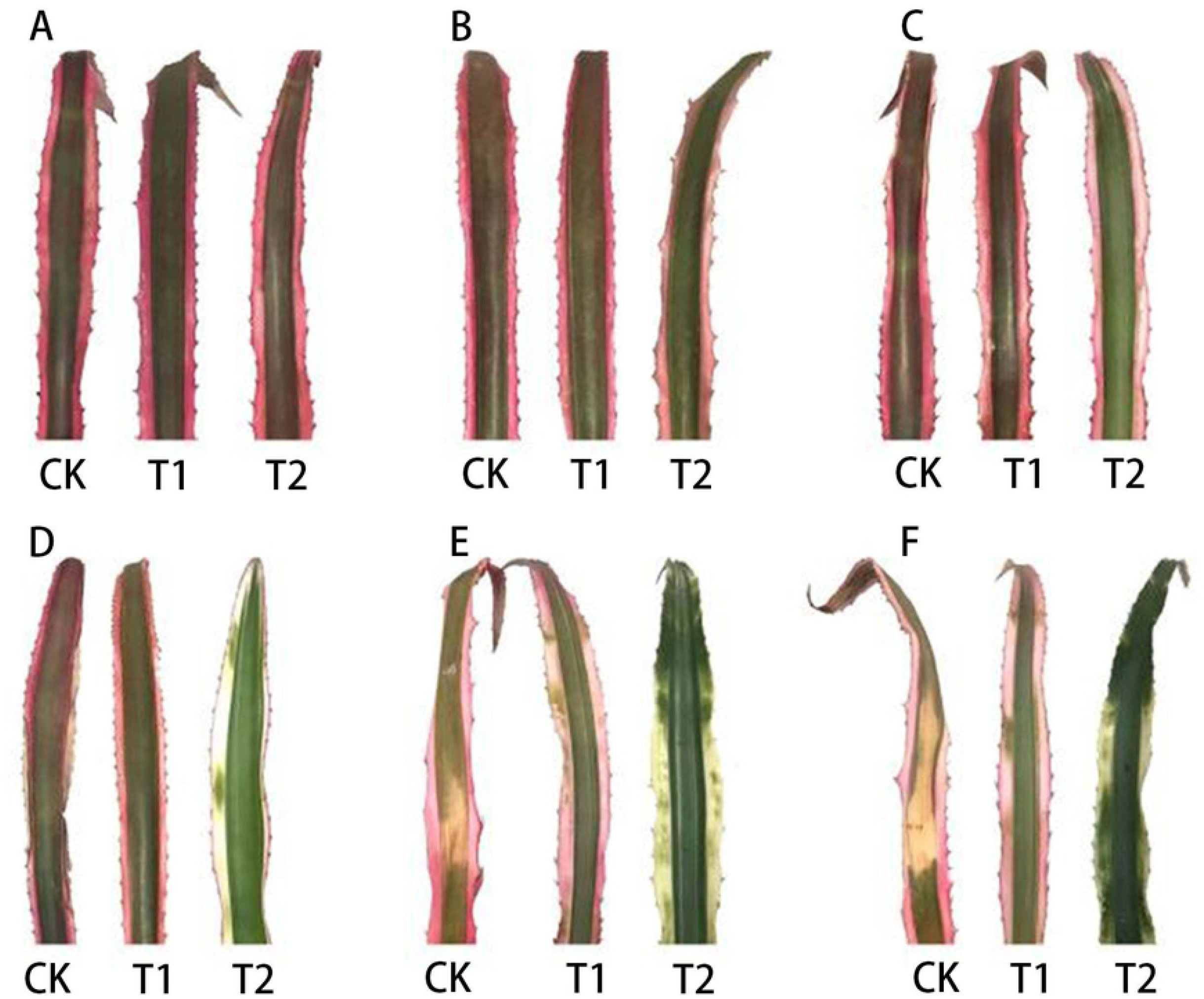
Visual foliage color changes of *Ananas comosus var. bracteatus* leaves under different light intensity. (CK) full light, (T1) 50% shade, (T2) 75% shade, (A) day 0, (B) day 15, (C) day 30, (D) day 45, (E) day 60, (F) day 75.

In order to analyze the color change of the leaves furtherly, the leaf color parameters (L*, a*, b*) were detected. For treatment T1 and T2, value a* of the edge of leaves gradually decreased. L* (lightness) of the green part of leaves increased slightly as shade increased for all treatments (Fig 3A,B), indicating that the margin of leaves had a gradual transition from red to green for T1 and T2 and the green part of leaves became lighter for all treatments during our experiment (Fig 2A-F). The value a* and b* of the margin of leaves decreased faster than the green part. The chimeric leaves of three treatments kept red in the first 30 days and then lost the red color apparently. As light intensity decreased more and the shading time lasted longer, the leaf lost red color quickly. Under 75 days of 75% shading treatment, the leaf margin color moved from red to green. Shading influenced the leaf margin color apparently by decreasing the red color and increasing the green color [11]. The center green part of the chimeric leaves maintained green, and the leaf color did not change a lot along the treatment.

**Fig 3.**
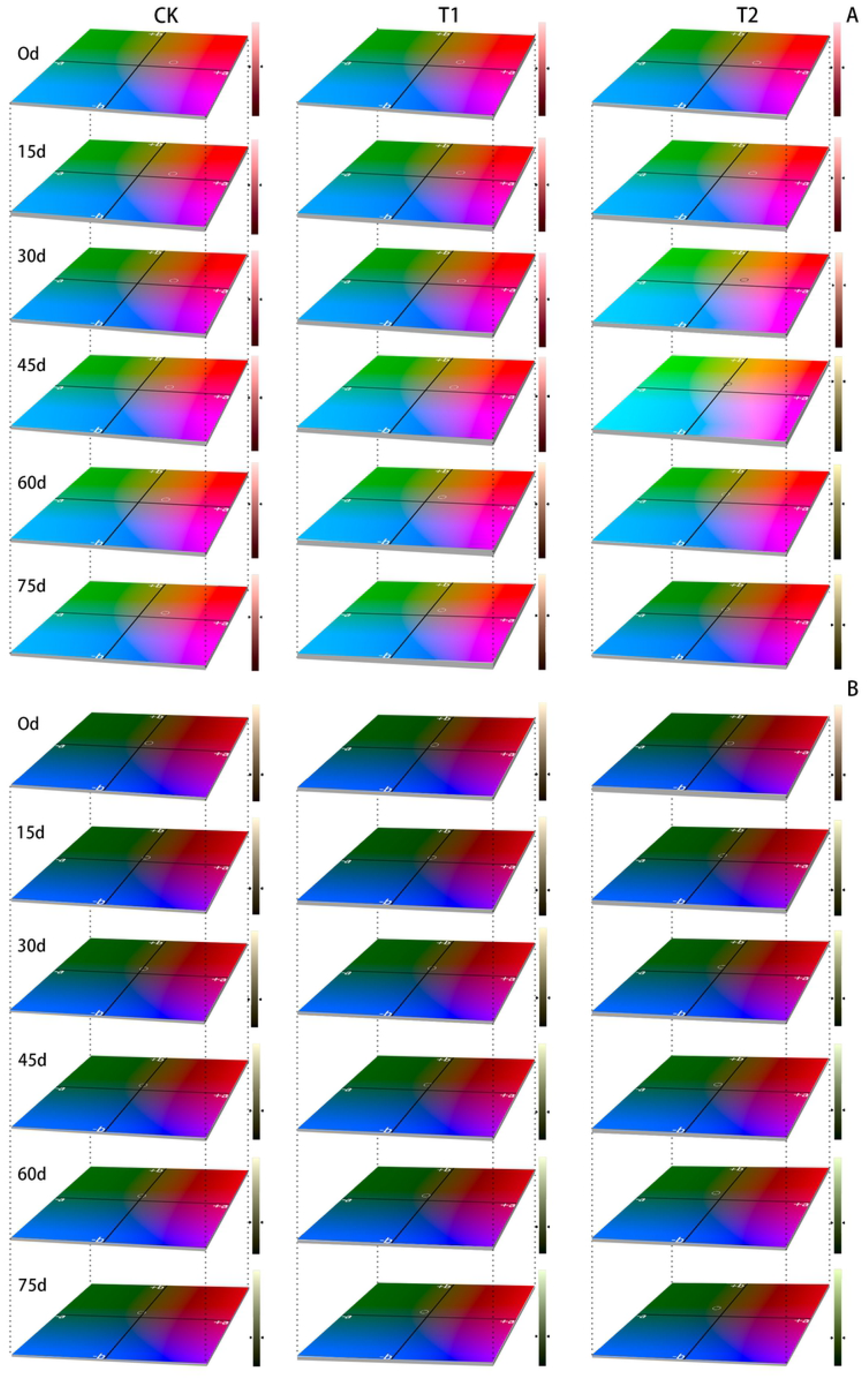
Changes in leaf color parameters of the green center part (A) and red margin part (B) of leaves under different light intensity. The circle in the color plate indicated the color measured.

### Pigment content changes in *Ananas comosus var. bracteatus* chimeric leaves

The leaf color is directly related to the pigments content of the leaves. In order to reveal the material basis of the leaf color changes, the pigment content of the margin and center part of chimeric leaves of *Ananas comosus var. bracteatus* were analyzed.

### Pigments changes in the center green part of the chimeric leaves

Shading had significant influences on the accumulation of chlorophyll, carotenoid and anthocyanin in the green center part of *Ananas comosus var. bracteatus* leaves (Fig 4A-C). The chlorophyll content of center part of chimeric leaves under full light and 50% shading treatment kept decreasing along 75 days. The chlorophyll content under full light treatment decreased faster than that under 50%. This finding may be related to a greater degradation caused by high solar radiation and leaf temperature under full light. Additionally, it was reported that being exposed to high light intensities for a long time may harm the photosynthetic apparatus [35]. Chlorophyll was usually synthesized and photo-oxidized in the presence of light. So the content of chlorophyll was greatly affected by shading [4,36]. Moreover, the degree of photosynthetic damage increased with the time of exposure to high solar radiation, and severe photo-inhibition was followed by leaf death [32]. It is also important to point out that there is a markable increase in leaf chlorophyll content under 75% shade for 45 days, which demonstrated that plant’s ability to develop mechanisms of light harvest and greater plasticity to face light deficit conditions by enhancing chlorophyll content [37]. The carotenoid content decreased apparently under full light and 50% shading treatment during the 75 days. But under 75% shading it decreased during the first 45 days and then increased apparently from 45 to 75 days (Fig. 4B). Increased carotenoids content under shading can enhance light absorption and transfer to chlorophyll for photosynthesis [38]. During the experiment time, anthocyanins content of the green center part of the leaves were all decreased consequently. When the light intensity decreased, the content of anthocyanin decreased more. The highest anthocyanin content was observed under full light (Fig 4C). Previous study showed that anthocyanins have the ability to protect chlorophyll pigments from damage in high solar radiation conditions [39].

**Fig 4.**
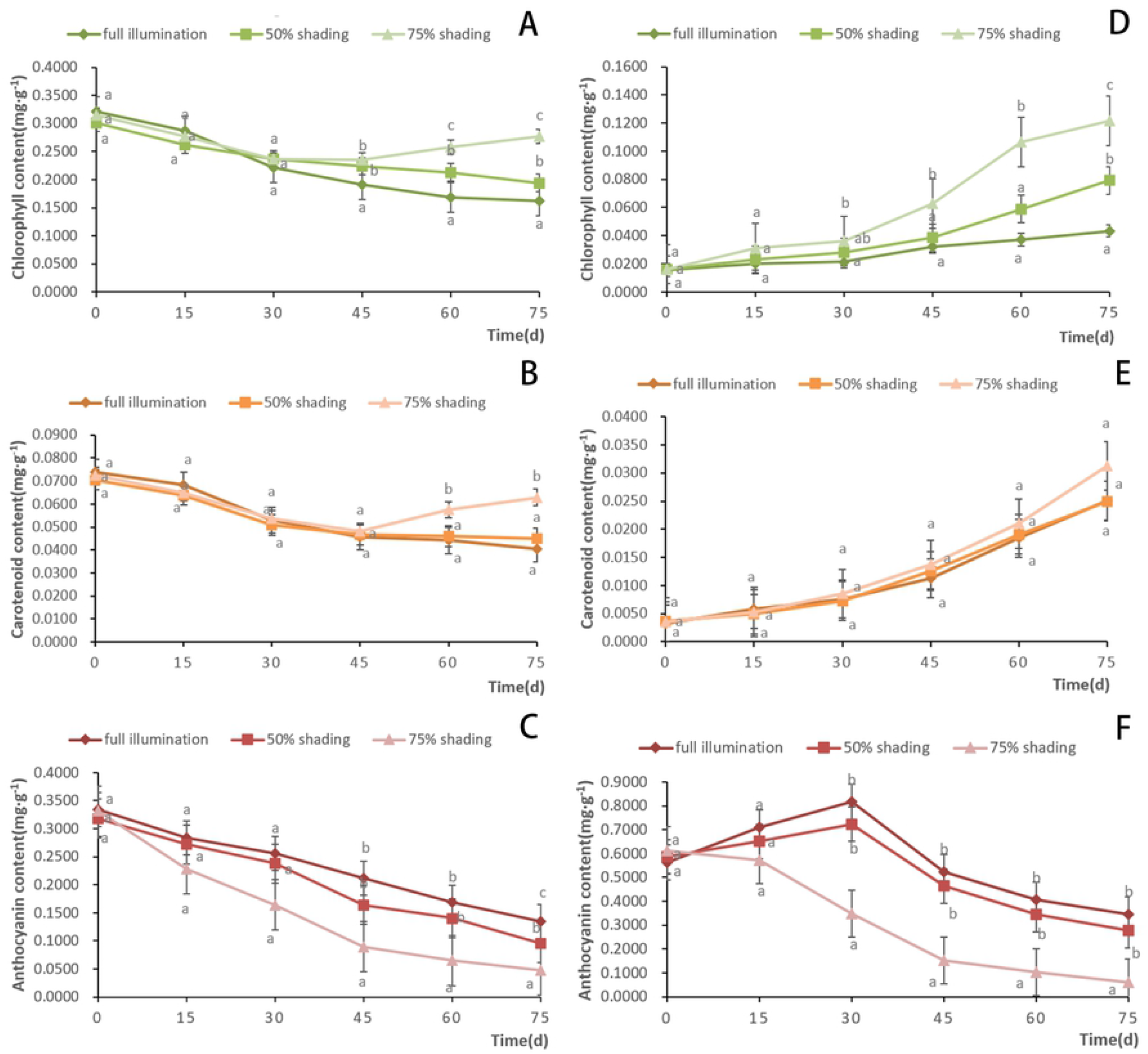
The changes of chlorophyll (A), carotenoid (B) and anthocyanin (C) content in green center part of *Ananas comosus var. bracteatus* leaves and the changes of chlorophyll (D), carotenoid (E) and anthocyanin (F) content in the red margin part under different shading treatments

### Pigment changes in the red margin part of the chimeric leaves

During the treatment period, the chlorophyll contents of the red margin part of chimeric leaves were increased under three treatments. There are apparent differences in the increasing chlorophyll content among the three treatments. Under 75% shading, the chlorophyll content was the highest and increased most rapidly, especially after 45 days of treatment (Fig 4D). Along the treatment procedure, the carotenoid content of marginal part of the leaves increased consequently under three treatments. But there are no significant differences in carotenoid content among the three treatments. (Fig 4E). However, shading can fast the decrease of anthocyanin content in the red margin part of leaves (Fig 4F). The content of anthocyanin decreased more when the shading intensity and time increased. At the first 30 days of full light and 50% shading, the anthocyanin content increased, and then decreased quickly during 30 days to 75 days. Under 75% shading, the anthocyanin content decreased consequently along the 75 days. It has been shown that light exposure was a prerequisite for significant anthocyanin synthesis, and a high level of solar radiation promoted the anthocyanin synthesis [9,10]. In addition, anthocyanin in epidermal tissue can provide some protection from UV solar radiation [40]. In order to maintain plants growth, chlorophyll gradually accumulated in the red part while the anthocyanin content decreased after day 30.

### Proportion of pigments in *Ananas comosus var. bracteatus* chimeric leaves

The leaf color is determined by the content and relative proportion of the chlorophyll, carotenoid and anthocyanin. In the green center part of the chimeric leaves, chlorophyll and anthocyanin were the dominant pigments sharing about 45% of the total pigments amount respectively during the first 15 days, so the center part of the leaves showed dark green. This is the result of the interaction between the reflectance of green chlorophyll and red anthocyanin. During 30 to 75 days, the proportion of chlorophyll increased apparently. The increase of chlorophyll proportion is more apparent with the increase of shading intensity. After 75 days of 75% shading, the proportion of chlorophyll increased to about 70% and the proportion of anthocyanin decreased to about 10%. The proportion of carotenoid increased slightly along the shading time and shading intensity (Fig. 5A). According to the increase of chlorophyll and decrease of anthocyanin, the color of the center part of the chimeric leaves changed from dark green to green.

**Fig 5.**
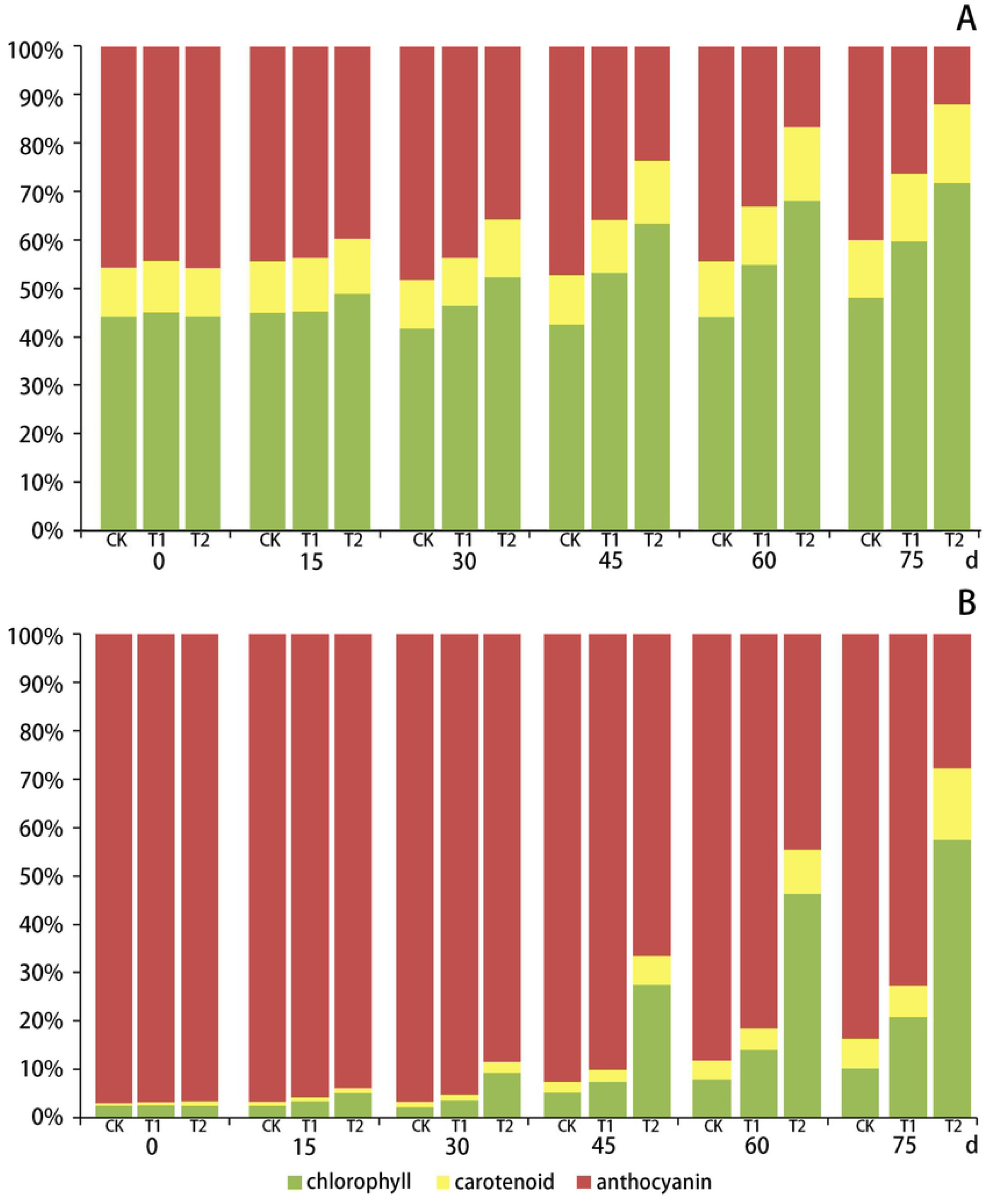
The changes of relative proportion of chlorophyll, carotenoid and anthocyanin in the green center part (A) and red margin part (B) of *Ananas comosus var. bracteatus* chimeric leaves under different light intensity.

In the red margin part of the chimeric leaves, anthocyanin was the dominant pigment, then chlorophyll and carotenoid. At the first 30 days, anthocyanin accounted for more than 90% of the total pigment amount. After 30 days of treatment, the proportion of chlorophyll and carotenoid increased and that of anthocyanin decreased (Fig. 5B). As the shading intensity and time increased, the chlorophyll proportion increased more and the anthocyanin proportion decreased more. Under full light treatment for 75 days, the proportion of anthocyanin content remained more than 80% of the total pigment content, while under 75 days of 75% shading treatment the proportion of anthocyanin decreased to 28%. Under 50 shading for 75 days, the anthocyanin proportion was more than 70%. The changes in the pigments relative proportion resulted in the morphology changes we observed. It is notable that the proportion of carotenoid in the red margin of the leaves increased more apparently than that in the green center part of leaves, especially under 75% of shading treatment. There are different pigment ratios in plants grown under different light conditions, and the ratios of pigments can reflect the light adaptability of plants. Plants grown under high light tend to produce more anthocyanin, which enhances the ornamental character of plants [9,12,40]. However, plants tend to produce more chlorophyll under low light condition to maintain sufficient photosynthetic antennas. This response allows the plant to capture the required light energy [8,10]. In addition, the different coloration between the center and marginal part of the chimeric leaves is the result of the different content and ratio between anthocyanin and chlorophyll. Shading can influence the pigment ratio in *Ananas comosus var. bracteatus,* which agrees with other studies [10,12].

### Pigment location and leaf structure changes of chimeric leaves

In order to reveal the distribution of pigment in the chimeric leaves and the effect of shading on the structure of the leaf tissues, anatomic observations of the leaves were carried out. The red chimeric leaves used in this experiment were consisted of red margin and green center tissues (Fig 6A). In the red margin part, no chloroplast was observed in the mesophyll cells, all the mesophyll cells are white (Fig 6B1,C1,E1). In the green center part of the leaf, large amounts of chloroplasts were observed in the mesophyll cells which gave it green color (Fig 6B1,D1,F1). Red cells containing anthocyanin were located in the upper and lower epidermis cells and the 1-2 cell layer beneath the epidermis of the leaf (Fig 6C1,D1). Anthocyanins were located in the vacuoles of the cells, which made the entire cell appear red. Under the full light for 75 days, the red cells with anthocyanin always existed in/under epidermis in both red margin and green center part of the leaf, but were less than that in leaves of 0 d (Fig 6A2,B2,C2,D2). Almost no green cells with chloroplasts were observed in the mesophyll cells of red leaf margin under full light (Fig 6B2,C2). However, a few green spots were observed in the red margin of the leaf under 50% shading for 75 days (Fig. 6A3) because some green cells with chloroplasts existed in the red margin (Fig 6B3,C3). Red cells of the leaf under 50% shading existed in/under the upper and lower epidermis of red margin and green center of the leaf, but less than that of the leaf of the 0 d (Fig 6B3,C3,D3). After treated with 75% shading for 75 d, the margin of the leaf lost red color and then turned green (Fig 6A4). Red cells were seldom observed in/under the epidermis of the leaf. However, green cells with chloroplasts apparently accumulated in the margin of the leaf (Fig 6B4,C4,D4). It is also important to point out that there was no coexistence of chlorophyll and anthocyanin in a cell.

**Fig 6.**
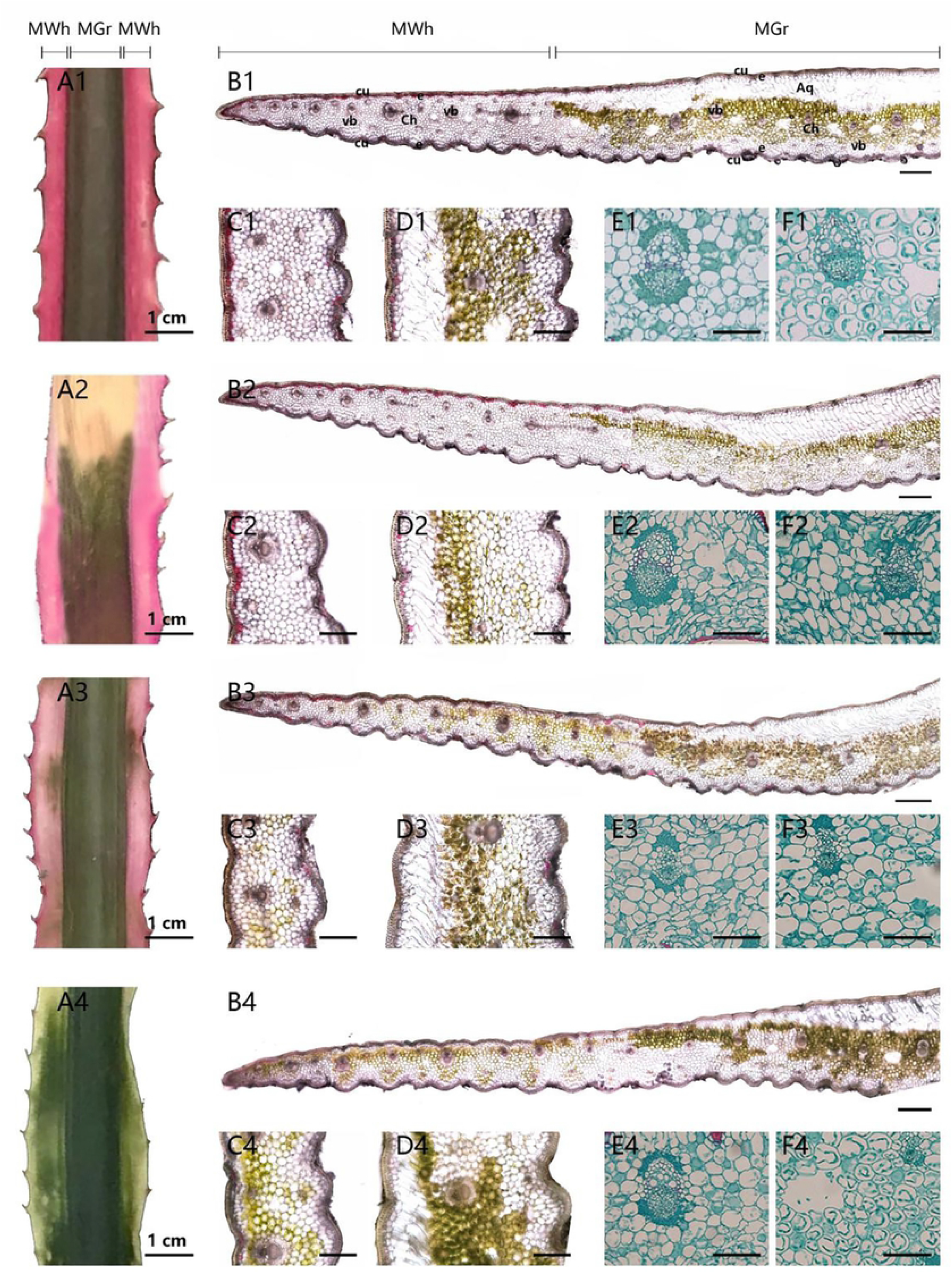
The pigment location and anatomic character of the green center and red margin part of the chimeric leaves. (A1-F1) The anatomic structure of the leaf at 0 d. (A2-F2) The anatomic structure of the leaf under full light (CK) for 75 days (A3-F3) The anatomic structure of the leaf under 50% shading (T1) for 75 days. (A4-F4) The anatomic structure of the leaf under 75% shading (T2) for 75 days. (A) Leaf sample (Scale bars=1cm). (B) Leaf cross-section (Scale bars=400μm). (C) The distribution of pigment in red margin tissues (Scale bars=200μm). (D) The distribution of pigment in green center tissues (Scale bars =200μm). (E) The mesophyll parenchyma cell in the white margin part of the chimeric leaves (Scale bars =100μm). (F) The mesophyll parenchyma cell in the green center part of the chimeric leaves (Scale bars =100μm). MWh, white tissue; Mgr, green tissue; Cu, cuticle; e, epidermis; Ch, mesophyll parenchyma; Aq, water storage parenchyma; vb, vascular bundle.

Light intensity has been considered an important factor for the photosynthetic mechanism. However, plants absorb more light than they need to saturate photosynthesis and cause the damage of the photosynthetic machinery. With time of exposure to high light conditions, the degree of damage will increase and leaf death will occur [31]. In our present study, full light was much for *Ananas comosus var. bracteatus*. As is already known, light intensity promotes structural changes in leaf tissue. The mesophyll cells in the leaf margin and green center tissues of the leaves under full light were reduced in size, and some of them were wrinkled in transverse section (Fig 6 E2,F2). It could be that excessive light and high leaf temperature resulted in water losing of the leaf cells. However, the mesophyll cells under 75% shading were oval and tense (Fig.6 E4, F4). Under the cuticle of leaf center green part presented large parenchyma cells, which were water storage cells, an important structural feature of *Ananas comosus var. bracteatus* as a drought-enduring plant adapting to the drought environment. This is an important structure of the green center part of chimeric leaves.

## Conclusions

In this study, our results can be concluded that light intensity affects the leaf color, color parameters, pigments accumulation, distribution and ratio, mesophyll cells, and chloroplasts of the marginal and central parts of chimeric leaves of *Ananas comosus var. bracteatus.* 75% shading boosted the decrease of anthocyanin content, increase of chlorophyll content, and the number of chloroplasts in the mesophyll cells, and the foliage transited its color to green reducing the ornamental value. However, high irradiance promoted the accumulation of anthocyanin in the mesophyll cells, relative proportion of anthocyanin content and value L* (lightness) of the margin part of chimeric leaves. The plants submitted to full light and 50% shade could maintain red and had a higher ornamental value. In addition, long exposure to full light contributed to light damage or even leaf death and the occurrence of wrinkled and smaller mesophyll cells in *Ananas comosus var. bracteatus,* while 50% shade could prevent the light damage caused by full light. In order to balance its growth and ornamental value, 50% shading is a good cultivating condition for *Ananas comosus var. bracteatus* in summer.

## Acknowledgments

We are thankful to Sichuan Agricultural University, China, for providing support for the present investigation.

## Author contributions

**Conceptualization:** Jun Ma

**Data curation:** Wei Yang, Ma Jun

**Formal analysis:** Wei Yang, Yuke Lin

**Funding Acquisition:** Jun Ma

**Investigation:** Wei Yang, Yuke Lin, Yanbin Xue and Meiqin Mao

**Methodology:** Wei Yang, Xuzixing Zhou, Hu Hao and Lijun Feng

**Project administration:** Jun Ma

**Resources:** Jun Ma

**Software:** Wei Yang, Jiawen Liu and Huiling Zhang

**Supervision:** Jun Ma

**Validation:** Jun Ma

**Visualization:** Wei Yang, JiaHeng Luo

**Writing-original draft:** Wei Yang

**Writing-review & editing:** Jun Ma, Wei Yang

## Reference

1. Yang Y, Chen X, Xu B, Li Y, Ma Y, Wang G. Phenotype and transcriptome analysis reveals chloroplast development and pigment biosynthesis together influenced the leaf color formation in mutants of *Anthurium andraeanum* ‘Sonate.’ Front Plant Sci. 2015;6: 1–16. doi:10.3389/fpls.2015.00139

2. Wang Y, Liu S, Tian X, Fu Y, Jiang X, Li Y, et al. Influence of light intensity on chloroplast development and pigment accumulation in the wild-type and etiolated mutant plants of *Anthurium andraeanum* ‘Sonate.’ Plant Signal Behav. 2018;13. doi:10.1080/15592324.2018.1482174

3. Zhao MH, Li X, Zhang XX, Zhang H, Zhao XY. Mutation mechanism of leaf color in plants: A review. Forests. 2020;11: 1–19. doi:10.3390/F11080851

4. Dai Y, Shen Z, Liu Y, Wang L, Hannaway D, Lu H. Effects of shade treatments on the photosynthetic capacity, chlorophyll fluorescence, and chlorophyll content of *Tetrastigma hemsleyanum* Diels et Gilg. Environ Exp Bot. 2009;65: 177–182. doi:10.1016/j.envexpbot.2008.12.008

5. Fischer R, Turner NC. Plant Productivity in the Arid and Semiarid Zones. Ann RevPlant Physiol. 1978;29: 277–317.

6. Surpin M, Larkin RM, Chory J. Signal transduction between the chloroplast and the nucleus. Plant Cell. 2002;14: 327–338. doi:10.1105/tpc.010446

7. Norcini JG, Andersen PC, Knox GW. Light Intensity Influences Leaf Physiology and Plant Growth Characteristics of *Photinia × fraseri*. J Am Soc Hortic Sci. 2019;116: 1046–1051. doi:10.21273/jashs.116.6.1046

8. Kong DX, Li YQ, Wang ML, Bai M, Zou R, Tang H, et al. Effects of light intensity on leaf photosynthetic characteristics, chloroplast structure, and alkaloid content of *Mahonia bodinieri* (Gagnep.) Laferr. Acta Physiol Plant. 2016;38: 1–15. doi:10.1007/s11738-016-2147-1

9. Hughes NM, Neufeld HS, Burkey KO. Functional role of anthocyanins in high-light winter leaves of the evergreen herb *Galax urceolata*. New Phytol. 2005;168: 575–587. doi:10.1111/j.1469-8137.2005.01546.x

10. Palaniswamy UR. Effect of light intensity on the pigment composition and oxalic acid concentrations in *kalamegh* (*Andrographis paniculata*) Leaf. Acta Hortic. 2005;680: 109–114. doi:10.17660/ActaHortic.2005.680.15

11. Brand MH. Shade influences plant growth, leaf color, and chlorophyll content of Kalmia latifolia L. Cultivars. HortScience. 1997;32: 206–208. doi:10.21273/hortsci.32.2.206

12. Nguyen P, Cin VD. The role of light on foliage colour development in coleus (*Solenostemon scutellarioides* (L.) Codd). Plant Physiol Biochem. 2009;47: 934–945. doi:10.1016/j.plaphy.2009.06.006

13. Shi MZ, Xie DY. Features of anthocyanin biosynthesis in pap1-D and wild-type Arabidopsis thaliana plants grown in different light intensity and culture media conditions. Planta. 2010;231: 1385–1400. doi:10.1007/s00425-010-1142-9

14. Pereira TAR, da Silva LC, Azevedo AA, Francino DMT, Coser TDS, Pereira JD. Leaf morpho-anatomical variations in *Billbergia elegans* and *Neoregelia mucugensis* (Bromeliaceae) exposed to low and high solar radiation. Botany. 2013;91: 327–334. doi:10.1139/cjb-2012-0276

15. Wu J, Li J, Su Y, He Q, Wang J, Qiu Q, et al. A morphophysiological analysis of the effects of drought and shade on *Catalpa bungei* plantlets. Acta Physiol Plant. 2017;39. doi:10.1007/s11738-017-2380-2

16. Trojak M, Skowron E. Role of anthocyanins in high-light stress response. World Sci News. 2017;81: 150–168. Available: www.worldscientificnews.com

17. Coste S, Roggy JC, Imbert P, Born C, Bonal D, Dreyer E. Leaf photosynthetic traits of 14 tropical rain forest species in relation to leaf nitrogen concentration and shade tolerance. Tree Physiol. 2005;25: 1127–1137. doi:10.1093/treephys/25.9.1127

18. Valladares F, Niinemets Ü. Shade tolerance, a key plant feature of complex nature and consequences. Annu Rev Ecol Evol Syst. 2008;39: 237–257. doi:10.1146/annurev.ecolsys.39.110707.173506

19. Xue Y, Ma J, He Y, Yu S, Lin Z, Xiong Y, et al. Comparative transcriptomic and proteomic analyses of the green and white parts of chimeric leaves in *Ananas comosus var. bracteatus*. PeerJ. 2019;7: e7261 doi:10.7717/peerj.7261

20. Mao M, Xue Y, He Y, Zhou X, Rafique F, Hu H, et al. Systematic identification and comparative analysis of lysine succinylation between the green and white parts of chimeric leaves of *Ananas comosus var. bracteatus*. BMC Genomics. 2020;21: 1–15. doi:10.1186/s12864-020-6750-6

21. Ma J, Kanakala S, He Y, Zhang J, Zhong X. Transcriptome sequence analysis of an ornamental plant, *Ananas comosus var. Bracteatus*, revealed the potential unigenes involved in terpenoid and phenylpropanoid biosynthesis. PLoS One. 2015;10. doi:10.1371/journal.pone.0119153

22. Li X, Kanakala S, He Y, Zhong X, Yu S, Li R, et al. Physiological characterization and comparative transcriptome analysis of white and green leaves of *Ananas comosus var. bracteatus*. PLoS One. 2017;12: 1–17. doi:10.1371/journal.pone.0169838

23. Xiong YY, Ma J, He YH, Lin Z, Li X, Yu SM, et al. High-throughput sequencing analysis revealed the regulation patterns of small RNAs on the development of *A. comosus var. bracteatus* leaves. Sci Rep. 2018;8: 1–11. doi:10.1038/s41598-018-20261-z

24. Lin Z, Xiong Y, Xue Y, Mao M, Xiang Y, He Y, et al. Screening and characterization of long noncoding RNAs involved in the albinism of *Ananas comosus var. Bracteatus* leaves. PLoS One. 2019;14: 1–21. doi:10.1371/journal.pone.0225602

25. Ren J, Liu Z, Chen W, Xu H, Feng H. Anthocyanin degrading and chlorophyll accumulation lead to the formation of bicolor leaf in ornamental *kale*. Int J Mol Sci. 2019;20: 603. doi:10.3390/ijms20030603

26. Bae CH, Abe T, Nagata N, Fukunishi N, Matsuyama T, Nakano T, et al. Characterization of a periclinal chimera variegated tobacco (*Nicotiana tabacum* L.). Plant Sci. 2000;151: 93–101. doi:10.1016/S0168-9452(99)00205-8

27. Wang N, Zhu T, Lu N, Wang Z, Yang G, Qu G, et al. Quantitative phosphoproteomic and physiological analyses provide insights into the formation of the variegated leaf in *Catalpa fargesii*. Int J Mol Sci. 2019;20. doi:10.3390/ijms20081895

28. Marcotrigiano M. Genetic mosaics and the analysis of leaf development. Int J Plant Sci. 2001;162: 513–525. doi:10.1086/320138

29. Holm G. Chlorophyll Mutations in Barley. Acta Agric Scand. 1954;4: 457–471. doi:10.1080/00015125409439955

30. Liu X, Pan Y, Liu C, Ding Y, Wang X, Cheng Z, et al. Cucumber fruit size and shape variations explored from the aspects of morphology, histology, and endogenous hormones. Plants. 2020;9: 1–17. doi:10.3390/plants9060772

31. Kull O. Acclimation of photosynthesis in canopies: Models and limitations. Oecologia. 2002;133: 267–279. doi:10.1007/s00442-002-1042-1

32. Ludlow MM, Björkman O. Paraheliotropic leaf movement in Siratro as a protective mechanism against drought-induced damage to primary photosynthetic reactions: damage by excessive light and heat. Planta. 1984;161: 505–518. doi:10.1007/BF00407082

33. Newman SE, Follett MW. Irrigation Frequency and Shading Influences on Water Relations and Growth of Container-Grown *Euonymus japonica* ‘Aureo-marginata.’ J Environ Hortic. 1988;6: 96–100. doi:10.24266/0738-2898-6.3.96

34. Andersen PC, Norcini JG, Knox GW. Influence of Irradiance on Leaf Physiology and Plant Growth Characteristics of *Rhododendron* × ‘Pink Ruffles’. J Am Soc Hortic Sci. 2019;116: 881–887. doi:10.21273/jashs.116.5.881

35. Ramalho JC, Pons TL, Groeneveld HW, Azinheira HG, Nunes MA. Photosynthetic acclimation to high light conditions in mature leaves of *Coffea arabica* L.: Role of xanthophylls, quenching mechanisms and nitrogen nutrition. Aust J Plant Physiol. 2000;27: 43–51. doi:10.1071/pp99013

36. De Carvalho Gonçalves JF, De Sousa Barreto DC, Dos Santos UM, Fernandes AV, Barbosa Sampaio PDT, Buckeridge MS. Growth, photosynthesis and stress indicators in young rosewood plants (*Aniba rosaeodora* Ducke) under different light intensities. Brazilian J Plant Physiol. 2005;17: 325–334. doi:10.1590/s1677-04202005000300007

37. Lei TT, Tabuchi R, Kitao M, Koike T. Functional relationship between chlorophyll content and leaf reflectance, and light-capturing efficiency of Japanese forest species. Physiol Plant. 1996;96: 411–418. doi:10.1111/j.1399-3054.1996.tb00452.x

38. Czeczuga B. Carotenoid contents in leaves grown under various light intensities. Biochem Syst Ecol. 1987;15: 523–527. doi:10.1016/0305-1978(87)90098-6

39. Steyn WJ, Wand SJE, Holcroft DM, Jacobs G. Anthocyanins in vegetative tissues: A proposed unified function in photoprotection. New Phytol. 2002;155: 349–361. doi:10.1046/j.1469-8137.2002.00482.x

40. Burger J, Edwards GE. Photosynthetic efficiency, and photodamage by UV and visible radiation, in red versus green leaf coleus varieties. Plant Cell Physiol. 1996;37: 395–399. doi:10.1093/oxfordjournals.pcp.a028959

